# Sixteen thousand grafts reveal limited scion genetic control of grapevine bench-grafting success

**DOI:** 10.64898/2026.09.11.750947

**Authors:** Jose Munoz, Adam Lovgren, Dario Cantu, Luis Diaz-Garcia

## Abstract

Nurseries produce tens of millions of bench-grafted grapevines (*Vitis vinifera* L.) each year, yet the genetics underlying grafting success rate remain poorly understood. Here, we dissected the genetic architecture of this trait in a scion mapping population. We grafted 138 progeny from a Riesling × Cabernet Sauvignon cross, along with both parents, onto 1103P, with 40 grafts per genotype in each of three separate field blocks and grafted on separate days (16,645 grafts). Success averaged 91.1 %. Scion genotype explained 19 % of the variance among blocks (entry-mean heritability 0.42); genotype × block variance exceeded genotype variance, and a few genotypes failed almost completely in a single block. Success was uncorrelated with yield of the mother vines. QTL mapping across six phenotype definitions and four haplotype-resolved parental genome assemblies detected one modest, reproducible QTL on chromosome 7 (Cabernet Sauvignon alleles; +3.9 percentage points; 11 to 15 % of variance) and suggestive regions on chromosomes 5 and 9. Most variation in grafting success within this elite *V. vinifera* cross was non-genetic and attributable to events affecting entire bundles. These results set expectations for effect sizes, replication, and process control in genetic studies of grafting success.

## Introduction

Nearly all commercial grapevines (*Vitis vinifera* L.) are propagated by bench grafting, which joins two genotypes through intrinsic wound-healing processes: callus proliferation at the interface, formation of a continuous cambium, and reconnection of vascular tissue. These processes, together with adventitious rooting of the rootstock and growth of the scion bud, determine whether a graft becomes a saleable vine (Habibi *et al*., 2022; Loupit, Brocard, *et al*., 2023). Large nurseries produce millions of grafted vines per year, so the proportion of successful grafts directly affects production costs and the availability of planting material. Success rates vary among cultivar combinations, across years, and among wood lots within the same combination; the causes of this variation are only partially understood (Villa-Llop *et al*., 2025). Anatomical, biochemical, and transcriptomic correlates of graft incompatibility have been described in fruit crops, including grapevine (Assunção *et al*., 2016; Assunção *et al*., 2019; Camboué *et al*., 2025; Chen *et al*., 2017; Ciobotari *et al*., 2010; Cookson *et al*., 2014; Ermel *et al*., 1999; Hamza *et al*., 2026; Loupit, Valls Fonayet, *et al*., 2023; Loupit *et al*., 2025; Rehman *et al*., 2025).

By contrast, the quantitative genetics of grafting success remains largely unexplored. In apricot, graft incompatibility with a common rootstock segregates quantitatively and two QTL explain 14 to 16 % of the variance (Pina *et al*., 2021). In kiwifruit, where grafting is being adopted to combat vine decline syndrome, the causes of incompatibility remain largely unknown and only candidate markers and screening techniques have been proposed (Ashraf *et al*., 2025). In grapevine, genetic analyses of grafted material have focused on the rootstock side: root-related traits in rootstock populations are polygenic, with QTL explaining 3 to 14 % of variance (Blois *et al*., 2023; Morel *et al*., 2026). Whether scion genotype measurably contributes to nursery grafting success has not been tested. This question matters for breeding, because new scion cultivars must pass through the nursery pipeline, and for the design of any marker-based screening of propagation performance.

Estimating scion and rootstock genetic effects on grafting success is demanding because the phenotype of each genotype is a percentage, the success rate of a batch of grafts, and because many determinants of graft take, such as cane maturity and reserves, hydration, pathogen status, and callusing conditions, vary among wood lots and grafting days in ways that are difficult to measure (Villa-Llop *et al*., 2025). Distinguishing a genotype effect of a few percentage points from batch-to-batch noise requires many grafts per genotype and replication across independent batches. Here, we conducted a commercial-scale experiment with 16,645 grafts across three blocks grafted from different mother-vine blocks on different days (Figure 1), to quantify the genetic and non-genetic components of grafting success in a scion mapping population whose parents both have haplotype-resolved genome assemblies.

**Figure 1.**
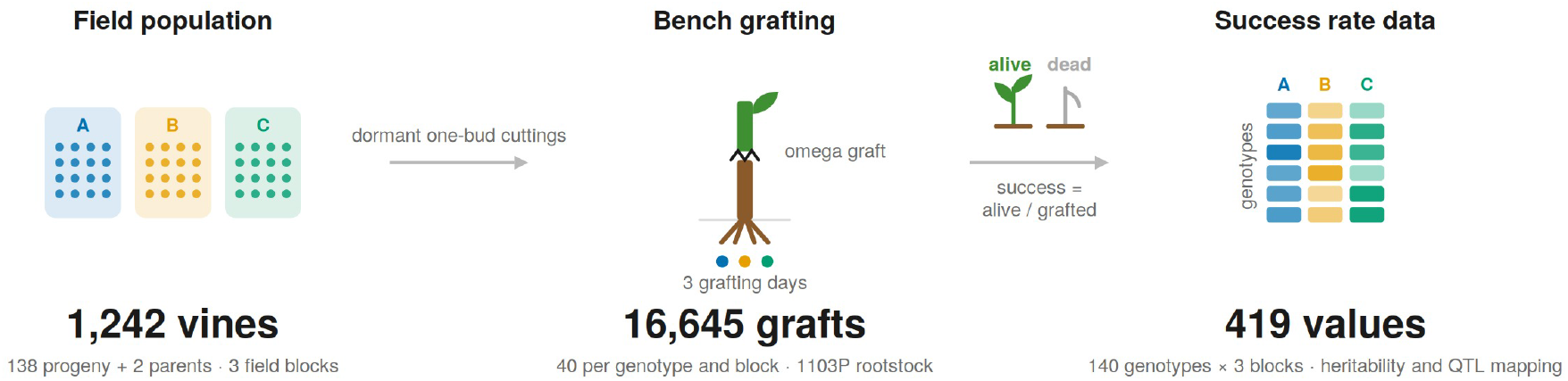
Experimental design. Dormant one-bud cuttings of 138 Riesling × Cabernet Sauvignon progeny and the two parents were collected separately from the three field blocks of the Oakville planting (three vines per genotype per block; 1,242 vines) and bench-grafted onto a common rootstock in three corresponding nursery blocks of 40 grafts per genotype, each grafted on its own day (16,645 grafts). After callusing and rooting, every vine was scored as alive or dead, and success was calculated as alive/grafted, giving 419 values (140 genotypes, three blocks); 134 genotyped progeny entered QTL mapping.

## Materials and methods

### 1. Plant material, grafting and phenotyping

The mapping population consists of 138 full-sibling F1 progeny from a cross between *Vitis vinifera* cv. Riesling (clone FPS 24/GM 110) and *V. vinifera* cv. Cabernet Sauvignon (clone FPS 08), made by M.A. Walker and C.P. Meredith in 1994. In 2017 the population was propagated, grafted onto 3309 Couderc, and transplanted to the UC Davis Department of Viticulture and Enology Experimental Station in Oakville, Napa County, CA (38°25′45.4″N; 122°24′36.4″W). The experiment design included three randomised blocks with three consecutive vines per genotype per block (138 progeny plus the two parents; 1,242 vines), 1.8 × 2.4 m vine and row spacing, modified vertical shoot positioning, two-bud spur pruning, drip irrigation, and a Cabernet Sauvignon border. The same planting has been used to study cluster architecture and yield components (Sharma *et al*., 2026), berry volatile composition (Lin, Massonnet, *et al*., 2026), wine aroma (Lin, Cantu, *et al*., 2026), and microbiome recruitment (Flörl *et al*., 2025).

Dormant one-bud scion cuttings were collected in February of 2025 separately from the three field blocks. Grafting was performed by Sunridge Nurseries (Bakersfield, CA, USA) from 27 to 30 May 2025 using omega bench grafting onto 1103P rootstock cuttings from a single source. Each field block was grafted as a single nursery block on one day (block A, 27 and 28 May; block B, 29 May; block C, 30 May), with 40 grafts per genotype per block (range 34 to 41; 16,645 grafts in total). Grafts were then callused and rooted under the nursery standard protocol. Vines were scored on [date] as alive (functional union, roots, and shoot growth) or dead, and success for each genotype in each block was calculated as alive/grafted (Figure 1).

### 2. Statistical analysis

Analyses were performed in R (ver. 4.3.3; R Foundation for Statistical Computing, Vienna, Austria). Variance components were estimated from the progeny (420 genotype × block observations) using three models: a linear mixed model on success, success ∼ block + (1|genotype), fitted by REML with lme4 (Bates *et al*., 2015); the same model on the empirical logit; and a binomial generalised linear mixed model on the vine counts with and without an observation-level random effect (OLRE) for overdispersion. Broad-sense heritability was calculated on a plot basis, H^2^ = V_G_/(V_G_ + V_E_), and on an entry-mean basis, H^2^ = V_G_/(V_G_ + V_E_/3), with parametric-bootstrap confidence intervals (500 samples). Genotype values for QTL mapping were the mean, median, and trimmed mean of the three blocks (the trimmed mean excluded a block more than 30 percentage points below the other two), the BLUPs from the linear and binomial models, and a rank-based normal score; single-block values were retained to assess consistency of QTL effects. Residuals were tested for association with field position, soil electrical conductivity, and elevation of the mother vines. Success was tested against yield of the mother vines, using BLUPs for clusters per vine, yield per vine, and cluster weight estimated from field phenotyping of the same planting in the 2023 and 2024 seasons (Sharma *et al*., 2026), for each season and for a two-season model; Spearman correlations were adjusted for multiple testing with the Benjamini–Hochberg procedure.

### 3. Genotyping and QTL mapping

Genotyping and linkage mapping of this population have been described previously, and the same genetic data have supported QTL mapping of cluster architecture and yield components, berry volatile composition, and microbiome recruitment (Flörl *et al*., 2025; Lin, Massonnet, *et al*., 2026; Sharma *et al*., 2026). Briefly, progeny and parents were genotyped by genotyping-by-sequencing (Glaubitz *et al*., 2014; Hyma *et al*., 2015), and reads were aligned to four haplotype-resolved parental genome assemblies: Cabernet Sauvignon FPS 08 v1.1 haplotypes 1 and 2 (Massonnet *et al*., 2020) and Riesling FPS 24 v1.1 haplotypes 1 and 2 (Lin, Massonnet, *et al*., 2026). For each reference, filtered SNPs were phased into parental haplotype calls and assembled into a Cabernet Sauvignon and a Riesling pseudo-testcross map (852 to 982 marker bins; 1,219 to 1,515 cM) with BatchMap (Schiffthaler *et al*., 2017), and composite maps combining the two parental maps were produced with LPmerge (Endelman & Plomion, 2014). Twelve cross objects were built with R/qtl (Broman *et al*., 2003) for the 134 genotyped progeny: eight parental maps coded as backcrosses and four composite maps coded as four-way crosses. For each cross object and phenotype, Haley-Knott interval mapping was run with 1,000 permutations for 5 %, 10 %, and 20 % genome-wide thresholds; non-parametric interval mapping (Kruglyak & Lander, 1995), which is robust to skewed distributions, was run for the mean and median (1,000 permutations); and composite interval mapping (three marker covariates, 10-cM window) was run on the parental maps for the BLUP and median (500 permutations). Support intervals are 1.5-LOD intervals expanded to flanking markers. Allele effects at peak markers were compared by Welch *t*-test.

## Results and discussion

### 1. Scion genotype explained a minority of the variance in grafting success

Across 419 genotype × block observations, success averaged 91.1 % (median 92.5 %) and was left-skewed (Figure 2A). Block means (90.6 to 91.4 %) did not differ significantly (*p* = 0.70). Cabernet Sauvignon (95.0 to 97.5 %) and Riesling (89.2 to 92.5 %) were close to the progeny mean, while progeny means ranged from 59.5 to 100 %, indicating transgressive segregation in both directions. Two progeny were consistently poor (DVIT002 and DVIT050), and four others failed almost completely in a single block while exceeding 78 % in the other two (Figure 2B). Correlations of genotype values between blocks were correspondingly low (*r* = 0.13 to 0.27; Figure 2C–E). Genotype was significant (likelihood-ratio χ^2^ = 14.1; *p* = 9 × 10^−5^) but explained only 19 % of block-level variance, yielding a broad-sense heritability of 0.42 (bootstrap 95 % CI 0.20 to 0.55) on an entry-mean basis (Table 1 and Supplementary Figure 1). Vine counts were strongly overdispersed relative to binomial sampling (dispersion ratio 1.84): with 40 grafts per bundle, binomial noise is small, and in the binomial mixed model the bundle-level variance was almost three times the genotype variance (Table 1). None of the nine yield × season combinations of the mother vines was correlated with success (|ρ| ≤ 0.08; FDR ≥ 0.90; Supplementary Table 1 and Supplementary Figure 2), and residuals showed no relationship with field position, soil conductivity, or elevation of the mother vines (Supplementary Figure 3).

**Figure 2.**
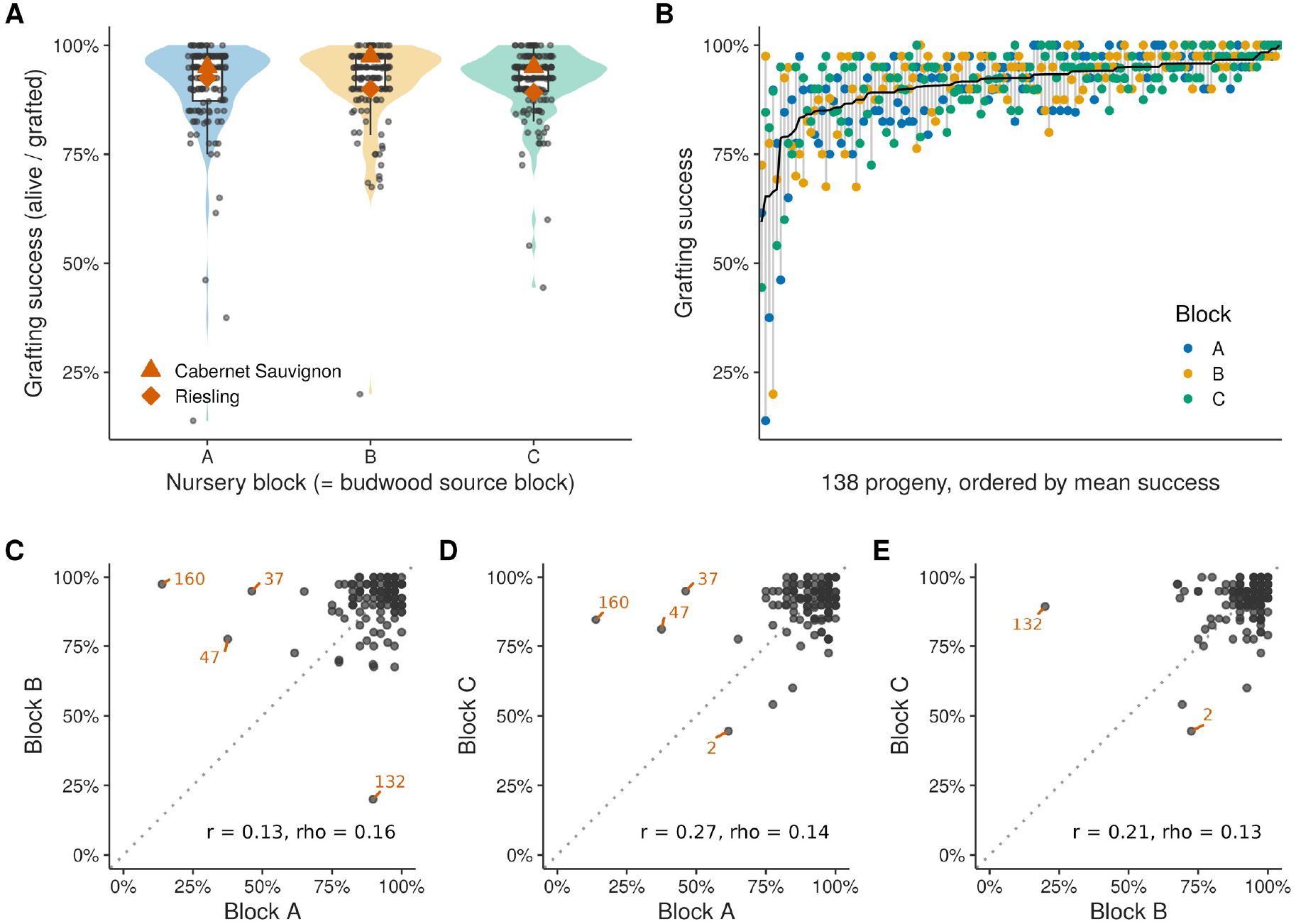
Grafting success in the Riesling × Cabernet Sauvignon progeny. Success (alive/grafted) of each genotype in each nursery block; parents are shown as orange symbols. (B) The 138 progeny ordered by mean success, with the three block values (coloured points) and the mean (line); grey segments span the block range. (C–E) Concordance of genotype values between blocks; genotypes with any block below 50 % are labelled. *r*, Pearson; ρ, Spearman.

**Table 1.** Variance components and broad-sense heritability of grafting success in 138 Riesling × Cabernet Sauvignon progeny grafted in three blocks of 40 vines.

| Model | VG | VE | H<sup>2</sup> (plot) | H<sup>2</sup> (entry mean) |
| --- | --- | --- | --- | --- |
| LMM rate | 0.0019 | 0.0080 | 0.19 | 0.42 |
| LMM elogit | 0.1710 | 0.7928 | 0.18 | 0.39 |
| Binomial GLMM | 0.4133 | – | – | – |
| Binomial GLMM + OLRE | 0.2105 | 0.5765 | – | – |

$V_G$ , genotype variance; $V_E$ , residual variance (observation-level variance for the binomial model with an observation-level random effect, OLRE); plot basis, single block; entry-mean basis, mean of three blocks.

The dominant source of variation was therefore not genotype but events affecting entire bundles. Because each genotype × block combination was one bundle of 40 cuttings collected from three mother vines and grafted on one day, many covariables are confounded within a bundle and most are difficult or impossible to measure retrospectively: cane maturity, diameter, and reserve status of the mother vines, water status at collection, hydration during storage, latent pathogen load, position in the callusing room, and handling by the grafting crew and day. Wood-lot effects of this kind are documented: success of a single Tempranillo/110 Richter combination varied with the field of origin of the rootstock canes, in association with anatomy, carbon isotope composition, and nitrogen status of the wood (Villa-Llop *et al*., 2025). Our data show they also operate on the scion side, at the scale of single bundles. Any study or screening programme for grafting success should treat the batch, not the graft, as the unit of replication: splitting genotypes into multiple bundles per day and randomising genotypes across grafting days would convert this variance into estimable error at no additional grafting cost.

### 2. QTL mapping detected one modest, reproducible region

Genome-wide 5 % thresholds were LOD 2.7 to 3.0 for the parental maps and 4.3 to 4.5 for the four-way maps, and LOD profiles from the four reference assemblies agreed closely (Figure 3). One region on chromosome 7 of the Cabernet Sauvignon map reached the 5 % level with the median phenotype (LOD 3.00 at 34.3 cM, peak marker at 10.96 Mb on haplotype 1; LOD 2.87 at 9.98 Mb on haplotype 2) and the 10 % level on both Riesling references, with LOD 2.2 to 2.7 for the other phenotype definitions (Table 2). Composite interval mapping strengthened this peak (LOD 4.41 and 5.28, above the 5 % thresholds, on the Cabernet Sauvignon references), and the region was recovered by non-parametric mapping and was the strongest signal on the four-way maps (Supplementary Table 2 and Supplementary Figure 4). Suggestive regions on chromosome 9 (Cabernet Sauvignon, 0 to 7 cM) and chromosome 5 (Riesling, 36 to 51 cM) were reproducible across references. Single-block phenotypes produced isolated peaks that did not appear with combined phenotypes, the pattern expected from bundle-level noise (Supplementary Figure 5). The 1.5-LOD support intervals of the three regions span 10 to 20 Mb (Supplementary Table 3).

**Figure 3.**
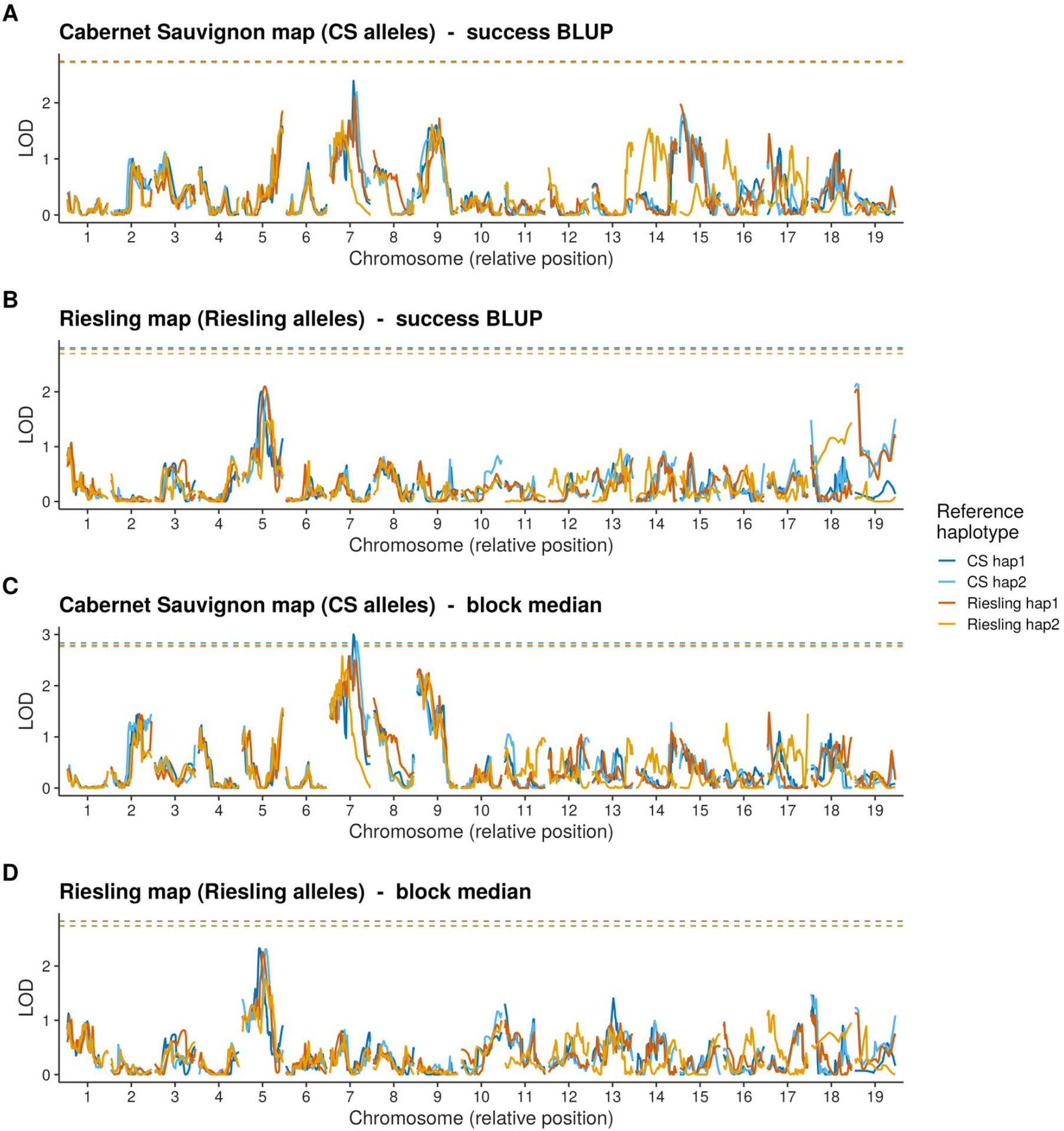
Genome scans on the parental pseudo-testcross maps. LOD profiles from Haley-Knott interval mapping on the four reference assemblies (colours), plotted on a relative chromosome scale; dashed lines are 5 % genome-wide thresholds from 1,000 permutations. (A and B) Success BLUP on the Cabernet Sauvignon and Riesling maps. (C and D) Block-median phenotype on the same maps.

**Table 2.** Chromosome peaks at or above the 20 % genome-wide level by Haley-Knott interval mapping.

| Chr | Map | Reference | Phenotype | cM | Mb | LOD | Thresholds | Level |
| --- | --- | --- | --- | --- | --- | --- | --- | --- |
| 5 | 4-way | RI-h1 | BLUP | 70.0 | 27.69<sup>a</sup> | 3.80 | 4.44 / 4.00 / 3.63 | 20% |
| 5 | 4-way | RI-h1 | Mean | 70.0 | 27.69<sup>a</sup> | 3.80 | 4.44 / 4.00 / 3.63 | 20% |
| 5 | RI | RI-h1 | BLUP (logit) | 41.0 | 11.09<sup>a</sup> | 2.33 | 3.03 / 2.62 / 2.26 | 20% |
| 5 | RI | RI-h1 | Median | 36.1 | 9.1 | 2.26 | 2.82 / 2.51 / 2.19 | 20% |
| 5 | RI | CS-h2 | Median | 51.0 | 8.11<sup>a</sup> | 2.31 | 2.83 / 2.52 / 2.22 | 20% |
| 5 | RI | CS-h1 | Median | 38.0 | 7.81<sup>a</sup> | 2.33 | 2.74 / 2.49 / 2.25 | 20% |
| 7 | 4-way | RI-h2 | Rank-normal | 23.9 | 7.2 | 3.78 | 4.43 / 4.05 / 3.60 | 20% |
| 7 | 4-way | RI-h2 | BLUP (logit) | 23.9 | 7.2 | 4.28 | 4.33 / 3.99 / 3.58 | 10% |
| 7 | 4-way | RI-h2 | BLUP | 23.9 | 7.2 | 4.26 | 4.31 / 3.96 / 3.55 | 10% |
| 7 | 4-way | RI-h2 | Trimmed mean | 23.9 | 7.2 | 3.79 | 4.23 / 3.88 / 3.48 | 20% |
| 7 | 4-way | RI-h2 | Median | 23.9 | 7.2 | 4.07 | 4.42 / 3.95 / 3.54 | 10% |
| 7 | 4-way | RI-h2 | Mean | 23.9 | 7.2 | 4.26 | 4.31 / 3.96 / 3.55 | 10% |
| 7 | CS | RI-h2 | Trimmed mean | 26.1 | 7.2 | 2.18 | 2.74 / 2.44 / 2.14 | 20% |
| 7 | CS | RI-h2 | Median | 26.1 | 7.2 | 2.58 | 2.78 / 2.47 / 2.22 | 10% |
| 7 | CS | RI-h1 | BLUP (logit) | 35.8 | 13.2 | 2.43 | 2.91 / 2.65 / 2.33 | 20% |
| 7 | CS | RI-h1 | Trimmed mean | 36.0 | 13.19<sup>a</sup> | 2.28 | 2.77 / 2.43 / 2.15 | 20% |
| 7 | CS | RI-h1 | Median | 28.3 | 10.2 | 2.58 | 2.79 / 2.53 / 2.26 | 10% |
| 7 | CS | CS-h2 | BLUP (logit) | 36.5 | 10.0 | 2.42 | 2.93 / 2.62 / 2.30 | 20% |
| 7 | CS | CS-h2 | Trimmed mean | 37.0 | 9.98<sup>a</sup> | 2.32 | 2.77 / 2.47 / 2.16 | 20% |
| 7 | CS | CS-h2 | Median | 36.5 | 10.0 | 2.87 | 2.76 / 2.49 / 2.23 | 5% |
| 7 | CS | CS-h1 | Rank-normal | 34.3 | 11.0 | 2.52 | 3.03 / 2.72 / 2.34 | 20% |
| 7 | CS | CS-h1 | BLUP (logit) | 34.3 | 11.0 | 2.70 | 2.94 / 2.62 / 2.29 | 10% |
| 7 | CS | CS-h1 | BLUP | 34.3 | 11.0 | 2.39 | 2.73 / 2.48 / 2.17 | 20% |
| 7 | CS | CS-h1 | Trimmed mean | 35.0 | 10.96<sup>a</sup> | 2.56 | 2.72 / 2.42 / 2.12 | 10% |
| 7 | CS | CS-h1 | Median | 34.3 | 11.0 | 3.00 | 2.84 / 2.50 / 2.22 | 5% |
| 7 | CS | CS-h1 | Mean | 34.3 | 11.0 | 2.39 | 2.73 / 2.48 / 2.17 | 20% |
| 9 | CS | RI-h2 | Median | 17.0 | 3.80<sup>a</sup> | 2.22 | 2.78 / 2.47 / 2.22 | 20% |
| 9 | CS | RI-h1 | Median | 4.0 | 2.48<sup>a</sup> | 2.32 | 2.79 / 2.53 / 2.26 | 20% |
| 9 | CS | CS-h2 | Median | 4.4 | 3.2 | 2.32 | 2.76 / 2.49 / 2.23 | 20% |

Map: CS, Cabernet Sauvignon pseudo-testcross map; RI, Riesling map; 4-way, composite map. Reference: CS-h1/h2, Cabernet Sauvignon haplotypes 1 and 2; RI-h1/h2, Riesling haplotypes 1 and 2. Mb, physical position of the peak marker on the given reference. Thresholds, genome-wide 5 % / 10 % / 20 % LOD from 1,000 permutations. Level, strongest genome-wide level reached. <sup>a</sup>LOD peak at an imputed position between genotyped markers; the value given is the physical position of the nearest genotyped marker (0.1 to 4.4 cM from the peak).

At the peak markers, the favourable Cabernet Sauvignon allele on chromosome 7 increased mean success from 88.9 to 92.8 % (median phenotype *p* = 4 × 10^−4^), with the same direction across all three independently grafted blocks; the chromosome 9 allele added 2.0 points and the Riesling chromosome 5 allele 3.5 points (Figure 4 and Supplementary Table 4). The chromosome 7 effect, 11 to 15 % of variance, was close to the upper limit of what this design could reliably detect, and even that estimate is likely inflated, because effect sizes at loci selected for significance tend to be overestimated in populations of this size. It is comparable to the graft-incompatibility QTL in apricot (14 to 16 % of variance; Pina *et al*., 2021) and to QTL for rooting and early growth in grapevine rootstock populations (Blois *et al*., 2023; Morel *et al*., 2026). The suggestive regions and the transgressive segregation indicate that both parents carry alleles that increase and decrease success; together these observations describe an oligogenic trait controlled by several loci of small effect. The support intervals are too wide for candidate gene analysis. Given the biology of union formation (Cookson *et al*., 2014; Loupit, Valls Fonayet, *et al*., 2023; Loupit *et al*., 2025), progeny contrasting at the chromosome 7 locus would be suitable material for asking at which stage the difference arises, for example by scoring callus development, interface metabolites, or xylem reconnection (Camboué *et al*., 2025).

**Figure 4.**
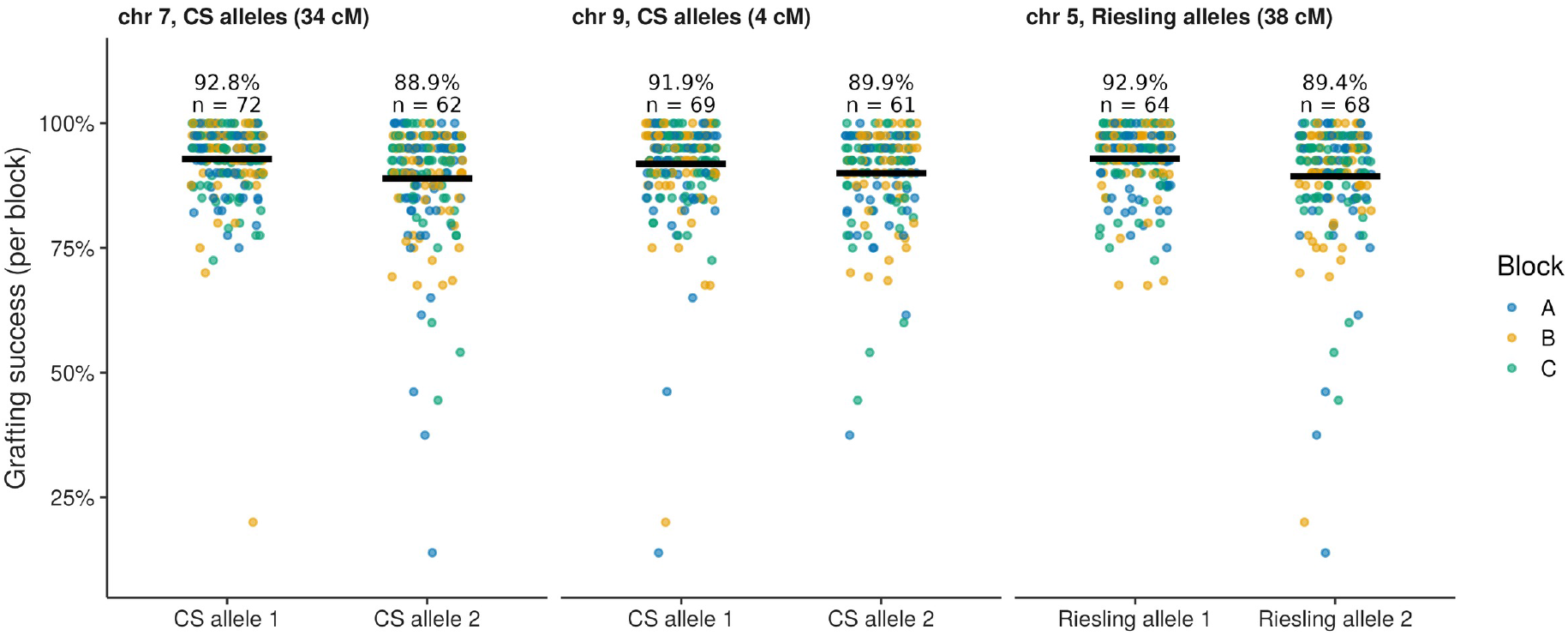
Allele effects at the three QTL regions. Success per block of progeny carrying either parental allele at the peak marker (points, coloured by block); bars are means; the mean of the genotype means and the number of progeny per class are given above each group.

### 3. Robustness across haplotype-resolved references, and limitations

Repeating every analysis on the four haplotype-resolved assemblies of the two parents provided an inexpensive robustness check. Maps, thresholds, and scans were computed four times against independent references, and the conclusions, including the negative ones, were the same on all four. The references are not interchangeable, because haplotype-specific variation, such as a structural variant present in only one haplotype, generates markers specific to that assembly during SNP calling. Agreement across references therefore shows that a signal rests on marker information shared among the haplotypes, whereas a signal detected on a single reference may reflect real haplotype-specific variation and warrants individual scrutiny. Here, the chromosome 5, 7, and 9 signals were recovered on all four references, and no QTL was confined to a single assembly.

Our experiment has three major limitations. First, the population samples one cross between two elite *V. vinifera* cultivars; wider crosses, interspecific scions, or combinations spanning species boundaries may segregate for larger incompatibility effects (Hamza *et al*., 2026; Rehman *et al*., 2025). Second, although common across QTL mapping studies in grapevine, the population size used in this experiment is relatively small, limiting our ability to detect QTLs, particularly those with small effect sizes. Finally, the trait was measured in a single nursery season on a single rootstock. Repetition across years is desirable but extremely expensive, because each additional year requires thousands of grafted vines, and testing several rootstocks multiplies the number of plants again. This is precisely why quantitative expectations from one large experiment are useful: scion genetic effects in elite material are of the order of a few percentage points per locus, heritability across independently grafted batches is about 0.4, and any experiment or association study aiming to detect such effects needs tens of grafts per genotype per batch, several independent batches, and explicit modelling of batch variance.

## Conclusion

In 16,645 bench grafts of a Riesling × Cabernet Sauvignon progeny on a common rootstock, scion genotype explained a fifth of the variance in grafting success among 40-graft bundles. One reproducible QTL on chromosome 7 and suggestive regions on chromosomes 5 and 9 each shifted success by two to four percentage points. Success was independent of mother-vine yield, and the largest source of variation was bundle-level events at the nursery for which no measured covariate accounted. Grafting success in elite *Vitis vinifera* scion material is a moderately heritable, oligogenic trait dominated by non-genetic batch effects, and its improvement is more likely to come from propagation process control than from scion selection.

## Acknowledgements

We thank Sunridge Nurseries for grafting, growing, and scoring the vines, and Guillermo Garcia-Zamora, Veronica Nunez, and Ana Gaspar for field and laboratory assistance. This work was supported by the California Rootstock Improvement Commission, the California Rootstock Research Foundation, the American Vineyard Foundation, and the USDA National Institute of Food and Agriculture Specialty Crop Research Initiative (2024-51181-43236 and 2026-67013-45985). Artificial intelligence (Claude, Anthropic) was used mostly for writing analysis and figure-generation code; all numbers and results obtained with AI-assisted code were rechecked by the authors.

## Author contributions

LDG: Conceptualization, Formal analysis, Writing - Original draft; AL : Conceptualization; JM: Data collection, Formal analysis, Writing - Review and editing; DC: Conceptualization, Writing - Review and editing.

## Conflicts of interest

The authors declare no conflicts of interest.

## Data availability

Raw genotyping-by-sequencing data for the mapping population are available as NCBI BioProject PRJNA1249123. Phenotypic data and analysis code are available from the corresponding author on request.

## Supplementary data

Supplementary Table 1. Spearman correlations between grafting success and mother-vine yield. Supplementary Table 2. Peaks from composite interval mapping and non-parametric interval mapping. Supplementary Table 3. 1.5-LOD support intervals of the three QTL regions. Supplementary Table 4. Allele effects at the peak markers. Supplementary Figure 1. Heritability of grafting success. Supplementary Figure 2. Grafting success and mother-vine yield. Supplementary Figure 3. Field position of the mother vines and model residuals. Supplementary Figure 4. Genome scans on the four-way composite maps. Supplementary Figure 5. Genome scans for all phenotype definitions.

## Notes

### Competing Interest Statement

The authors have declared no competing interest.

